# Genetic dissection of a *Solanum elaeagnifolium* tertiary genepool introgression associated with 4-*O*-caffeoylquinic acid accumulation in eggplant

**DOI:** 10.64898/2026.09.04.749343

**Authors:** Gloria Villanueva, Elena Rosa-Martínez, Diego A. Moreno, Pietro Gramazio, Virginia Baraja-Fonseca, Santiago Vilanova, Jaime Prohens, Mariola Plazas

## Abstract

Caffeoylquinic acids (CQAs) are the major phenolic compounds in eggplant (*Solanum melongena*) fruit, with 5-*O*-caffeoylquinic acid (5-CQA; chlorogenic acid) as the predominant isomer, while crop wild relatives may offer novel variation for diversifying these profiles. Previous work detected a distinct phenolic acid profile in *S. elaeagnifolium*, a tertiary genepool wild relative of eggplant, and in advanced backcross materials carrying introgressions from this species. This profile was characterized by a secondary chromatographic peak eluting after 5-CQA, but the identity of this compound and the associated chromosome 1 region remained unresolved. Here, we combined HPLC-MS-based metabolite identification, fine mapping, and comparative microsynteny analysis to characterize this phenotype. The secondary peak was identified as 4-*O*-caffeoylquinic acid (4-CQA). Fine mapping across three consecutive selfing generations, involving 612 screened individuals and 21 informative recombinants, reduced the associated interval from 2.753 Mb to 0.211 Mb, delimiting a region containing 30 annotated genes. The 4-CQA peak was detected in both homozygous and heterozygous introgression-derived materials, consistent with dominant control of the trait. Structural comparison of the fine-mapped interval revealed broad conservation between *S. melongena* and *S. elaeagnifolium*, but also identified a localized structurally divergent block potentially underlying the secondary 4-CQA peak. Within this block, the *S. elaeagnifolium* interval harbors acetylajmalan esterase-like/GDSL-type genes and a transcript-supported putative novel donor-specific gene. Overall, this study highlights *S. elaeagnifolium* introgressions as a promising source of phenolic diversification and providing markers to facilitate the development of eggplant lines with a more diverse CQA profile and potentially enhanced functional quality.

## 1. Introduction

Eggplant (*Solanum melongena* L.) is a widely consumed vegetable crop globally, valued for its economic importance, nutritional value, and health-promoting properties (Lyngdoh et al. 2025). Among the compounds contributing to eggplant fruit quality, phenolic acids are particularly relevant because they are major determinants of antioxidant capacity and are closely associated with the functional properties of the crop (Rosa-Martínez et al. 2023a; Lyu et al. 2025). Within this group, caffeoylquinic acids (CQAs) constitute the major phenolic fraction in eggplant fruit, with 5-*O*-caffeoylquinic acid (5-CQA), commonly referred to as chlorogenic acid (CGA), accounts for most of the total CQA content (Whitaker and Stommel 2003; Mennella et al. 2012). Because of their abundance and contribution to antioxidant properties, CQAs are important targets in breeding programmes aimed at improving the nutritional quality of eggplant fruit (Plazas et al. 2013b; Villanueva et al. 2024).

However, achieving significant improvements in these phenolic profiles depends on the availability of genetic variation, which is often limited within the cultivated gene pool (Kaushik et al. 2015; Meyer et al. 2015). In this respect, the use of crop wild relatives has contributed to broadening the variation of cultivated eggplant for agronomic, adaptive, and fruit quality traits (Gramazio et al. 2023), including compounds associated with nutritional value chlorogenic acid and other phenolic compounds (Rosa-Martínez et al. 2022; Martina et al. 2024). In eggplant, wild relatives have been reported to display broader hydroxycinnamic acid profiles than cultivated materials (CITA). These profiles include a wider diversity of caffeoylquinic acid-related compounds and derivatives, including positional isomers of 5-CQA such as 3-*O*-caffeoylquinic acid (3-CQA) and 4-*O*-caffeoylquinic acid (4-CQA) (Ma et al. 2010, 2011; Kaushik et al. 2017).

In parallel with the growing interest in improving fruit nutritional quality, eggplant has benefited from the development of genetic and genomic resources that facilitate the dissection of complex biochemical traits (Wei et al. 2020; Barchi et al. 2021; Gaccione et al. 2025). Linkage maps, QTL analyses, and high-throughput genotyping approaches have enabled the identification of genomic regions associated with fruit composition and other agronomically relevant traits, including chlorogenic acid and related metabolic compounds. In particular, some studies in eggplant have shown that candidate genes involved in chlorogenic acid biosynthesis are distributed across different chromosomes, while QTL associated with chlorogenic acid and other biochemical traits tend to cluster in some genomic regions (Gramazio et al. 2014; Toppino et al. 2016; Rosa-Martínez et al. 2023a; Gaccione et al. 2025). These advances provide a useful framework for studying the genetic control of phenolic compounds in eggplant and for linking metabolic variation with underlying genomic factors, particularly in populations carrying introgressions from wild relatives (Mangino et al. 2020; Rosa-Martínez et al. 2022; Villanueva et al. 2023).

Among eggplant crop wild relatives, *S. elaeagnifolium* Cav. is particularly interesting because it is an American wild species belonging to the tertiary genepool of eggplant and is highly divergent from cultivated eggplant and its closest relatives within the Eastern Hemisphere Spiny clade (Syfert et al. 2016; Aubriot and Knapp 2022). This evolutionary distance is reflected in reproductive barriers and structural differences between their genomes (Knapp et al. 2017). Despite these limitations, successful hybridization and backcrossing have enabled the development of eggplant advanced backcrosses introgressed with *S. elaeagnifolium* chromosome segments (García-Fortea et al. 2019; Villanueva et al. 2023). Overall, these breeding materials cover most (99.2%) of the genome of *S. elaeagnifolium* and constitute a valuable resource for uncovering novel allelic and metabolic variation absent from the domesticated gene pool. A previous study on these materials identified a distinct phenolic acid profile in *S. elaeagnifolium* and in some advanced backcrosses towards eggplant characterized by a secondary chromatographic peak eluting after the main chlorogenic acid (5-CQA) peak and associated this phenotype with a genomic region at the beginning of chromosome 1 (Villanueva et al. 2021). However, the identity of this secondary compound remained unresolved, and the genomic region associated with this phenotype was not further delimited or characterized.

In the present work, advanced backcross materials carrying introgressions from the wild relative *S. elaeagnifolium* in the *S. melongena* genetic background were selected for their distinct phenolic acid profile and characterized by the presence of a secondary peak eluting after the main 5-CQA peak. The objectives of this study were to identify the compound corresponding to this secondary CQA-related peak and to delimit more precisely and characterize the genomic region associated with this phenotype, thereby providing insight into the potential of these introgression materials as a source of novel metabolic variation for eggplant breeding. To address these objectives, we combined HPLC-MS-based metabolite identification, fine mapping, and comparative microsynteny analysis.

## 2. Material and Methods

### 2.1 Plant material and growth conditions

The plant material used in this study consisted of advanced backcross materials carrying introgressions of the wild relative *S. elaeagnifolium* (accession ELE2) in the *S. melongena* (accession MEL3) genetic background (Villanueva et al. 2023), as well as the recurrent MEL3 parent. Specifically, we selected materials carrying a broad genomic region on chromosome 1 that had previously been associated with a distinct phenolic acid profile, characterized by the presence of a secondary peak eluting after the main chlorogenic acid (5-CQA) peak (Villanueva et al. 2021).

From the available plant material, a single individual of the fifth backcross (BC5) of *S. elaeagnifolium* towards the recurrent parent *S. melongena* parent was selected to initiate the fine-mapping process. All genomic coordinates and interval boundaries reported in this study refer to the *S. melongena* reference genome ‘67/3’ v5 (GPE001970; Gaccione et al., 2025), hereafter referred to as the eggplant reference genome v5. This individual was chosen because it displayed the secondary CQA peak and carried the smallest available *S. elaeagnifolium* introgression covering the target region on chromosome 1, spanning 0.983-3.736 Mb according to the eggplant reference genome v5, which corresponds to 2.46% of the total chromosome length (112.110 Mb). The individual was heterozygous for both introgressions.

Three consecutive rounds of selfing and selection, one per year (2023 to 2025), were performed starting from the selected BC5 individual to fine-map the candidate region. In each generation, segregating populations of 222 (2023), 278 (2024), and 112 (2025) individuals were screened for recombination events using High-Resolution Melting (HRM), based on primer pairs designed to amplify polymorphic regions within the candidate interval on chromosome 1 (Table S1; see Section 2.3 for details). Based on this initial screening, subsets of 44 (2023), 28 (2024), and 24 (2025) putative recombinant individuals, respectively, were selected for high-throughput genotyping using Single Primer Enrichment Technology (SPET) (Barchi et al. 2019).

Seedlings were initially established in a climatic chamber under controlled photoperiod and temperature conditions of 16 h light (25 °C) and 8 h dark (18 °C). Following genotypic screening, selected individuals were transplanted into 15 L pots filled with coconut fiber substrate and cultivated on benches in a climate-controlled greenhouse at the Universitat Politècnica de València (Valencia, Spain), with temperatures ranging from 15 °C to 30 °C. Plants were supplied with water and nutrients through a drip fertigation system.

### 2.2 Caffeoylquinic acids extraction and analysis

The extraction of caffeoylquinic acids and related phenolic compounds from the flesh of eggplant fruits harvested at the commercial stage was performed following the protocol described by Plazas et al. (2014), with slight modifications. Freeze-dried samples (0.1 g) were extracted with 1.5 mL of methanol:water (80:20, v/v) containing 0.1% 2,3-tert-butyl-4-hydroxyanisole (BHT). The extracts were sonicated for one hour, centrifuged at 10,000 rpm for 3 minutes, and filtered through 0.2 µm PTFE membrane filters. Standard solutions of 5-*O*-caffeoylquinic acid (5-CQA; ≥95% purity, product no. C3878, Sigma-Aldrich, St. Louis, MO, USA) were prepared using the same protocol.

High-performance liquid chromatography (HPLC) analysis was performed using a Shimadzu SCL-40 system (Shimadzu Corporation, Kyoto, Japan) coupled to a diode array detector (DAD) and equipped with a Brisa “LC^2^” C18 column (3 μm; 150 × 4.6 mm; Teknokroma, Barcelona, Spain). The mobile phase consisted of water with 0.1% formic acid (solvent A) and methanol with 0.1% formic acid (solvent B), at a flow rate of 0.5 mL min^−1^. Detection was performed by absorbance at 325 nm.

In addition, a subset of extracts from nine advanced backcross materials was analysed by HPLCDAD-ESI-MSn to identify the additional chromatographic peak detected in some samples. Extracts were selected according to their HPLC profile and included samples showing the additional peak and representative samples showing only the main peak. Analyses were performed using an Agilent 1200 HPLC system (Agilent Technologies, Santa Clara, CA, USA) equipped with a photodiode array detector and coupled to a Bruker HCT-Ultra ion trap mass spectrometer (Bruker Daltoniks GmbH, Bremen, Germany) with an electrospray ionization (ESI) interface. Separation was carried out using the same Brisa “LC2” C18 column and mobile phases described above, at a flow rate of 0.8 mL min^−1^. Mass spectra were acquired in negative ion mode over an *m/z* range of 100-1200, and when necessary for compound annotation, MSn fragmentation data were obtained by collision-induced dissociation in the ion trap using He as collision gas. Peak identification was based on the combined evaluation of retention time, UV absorption spectrum, molecular ion signal, and fragmentation pattern, following the criteria described by Clifford et al. (2003).

### 2.3 Marker-assisted genotyping and selection

Genomic DNA was extracted from young leaves using the SILEX DNA extraction method (Vilanova et al. 2020). Primers targeting candidate genomic regions were designed based on polymorphic sites between the parental lines within the candidate interval on chromosome 1 (Table S1) and validated using the parental lines. Population screening was then performed by High-Resolution Melting (HRM) analysis using a LightCycler 480 Real-Time PCR system (Roche Diagnostics, Meylan, France). These HRM markers were used to identify recombinant individuals within the target region. Based on the HRM results, a subset of selected individuals was genotyped using high-throughput 5k Single Primer Enrichment Technology (SPET) (Barchi et al. 2019). The resulting Single Nucleotide Polymorphisms (SNPs) were filtered using Tassel software (version 5.2 Standalone) (Bradbury et al. 2007). Filtering criteria were adjusted to optimize each dataset, generally applying a minimum call rate of at least 95%, a Minor Allele Frequency (MAF) higher than 5%, and a maximum heterozygosity of approximately 70%. After filtering, the number of SNPs retained for analysis was as follows: 170 SNPs in the BC5 generation used to identify the individual that initiated the fine-mapping process, 100 SNPs in the first round of selfing and selection (2023), 117 SNPs in the second round (2024), and 42 SNPs in the third round (2025).

### 2.4 Whole-genome resequencing and sequence analysis

Whole-genome resequencing (WGRS) was performed on six genotypes, including the two parental lines and four selected recombinant BC5S2 individuals, corresponding to the second selfed generation after the fifth backcross. These recombinant plants were selected to enable a more detailed characterization of the target regions. High-quality DNA samples (with 260/280 and 260/230 ratios > 1.8) were submitted for the preparation of 150 bp paired-end libraries, followed by sequencing on the DNAseq platform at Beijing Genomics Institute (BGI Genomics, Hong Kong, China).

The raw sequencing data underwent quality control processing using fastq-mcf v.1.04.676 (Aronesty 2013), applying a quality threshold of 30 (-q 30) and a minimum sequence length of 50 bp (-l 50). The high-quality reads were aligned to the reference genome of eggplant v5 (Gaccione et al. 2025) using the BWA-MEM algorithm v. 0.7.17-r1188 with default settings (Li 2013). The resulting alignments were visually inspected in the Integrative Genomics Viewer (IGV) (Thorvaldsdóttir et al. 2013) to examine mapping patterns and genomic variations in specific regions of interest.

### 2.5 Candidate gene annotation and synteny analysis

Gene annotation for the *S. melongena* background in the candidate region was based on the eggplant reference genome v5 (Gaccione et al. 2025). Similarly, for *S. elaeagnifolium*, gene models were derived from a whole-genome annotation generated using the PASA pipeline (unpublished). This approach integrated de novo predictions from Helixer with full-length Iso-Seq transcript alignments (unpublished).

For the analysis of the fine-mapped candidate region, public *S. elaeagnifolium* transcriptomic data (SRR2038442, Tsaballa et al., 2015) were processed using RNA Galaxy workbench 2.0 (Fallmann et al. 2019). The resulting alignments were inspected in IGV to manually refine gene annotation and validate the orientation of novel genes. Orthologous relationships between *S. melongena* and *S. elaeagnifolium* were established based on sequence similarity, and the genomic organization of the region was visualized using the *gggenes* package in R (Wilkins 2023).

## 3. Results

### 3.1 Characterization of variation in the CQA profile

Representative HPLC chromatograms revealed clear differences in the caffeoylquinic acid (CQA) profile among *S. melongena, S. elaeagnifolium*, and the backcross-derived materials carrying either the full 2.75 Mb *S. elaeagnifolium* introgression or specific recombinant fragments of this region in chromosome 1 (Figure 1). The chromatographic profile of *S. melongena* was characterized by a single predominant peak at 21 min, whereas *S. elaeagnifolium* displayed a more complex pattern with multiple peaks across the chromatogram. Backcross-derived materials carrying the introgression on chromosome 1 exhibited a differential profile consisting of a predominant peak at 21 min together with a clearly resolved secondary peak at 22 min. This differential profile was used as the basis for subsequent peak identification and fine-mapping analyses.

**Figure 1.**
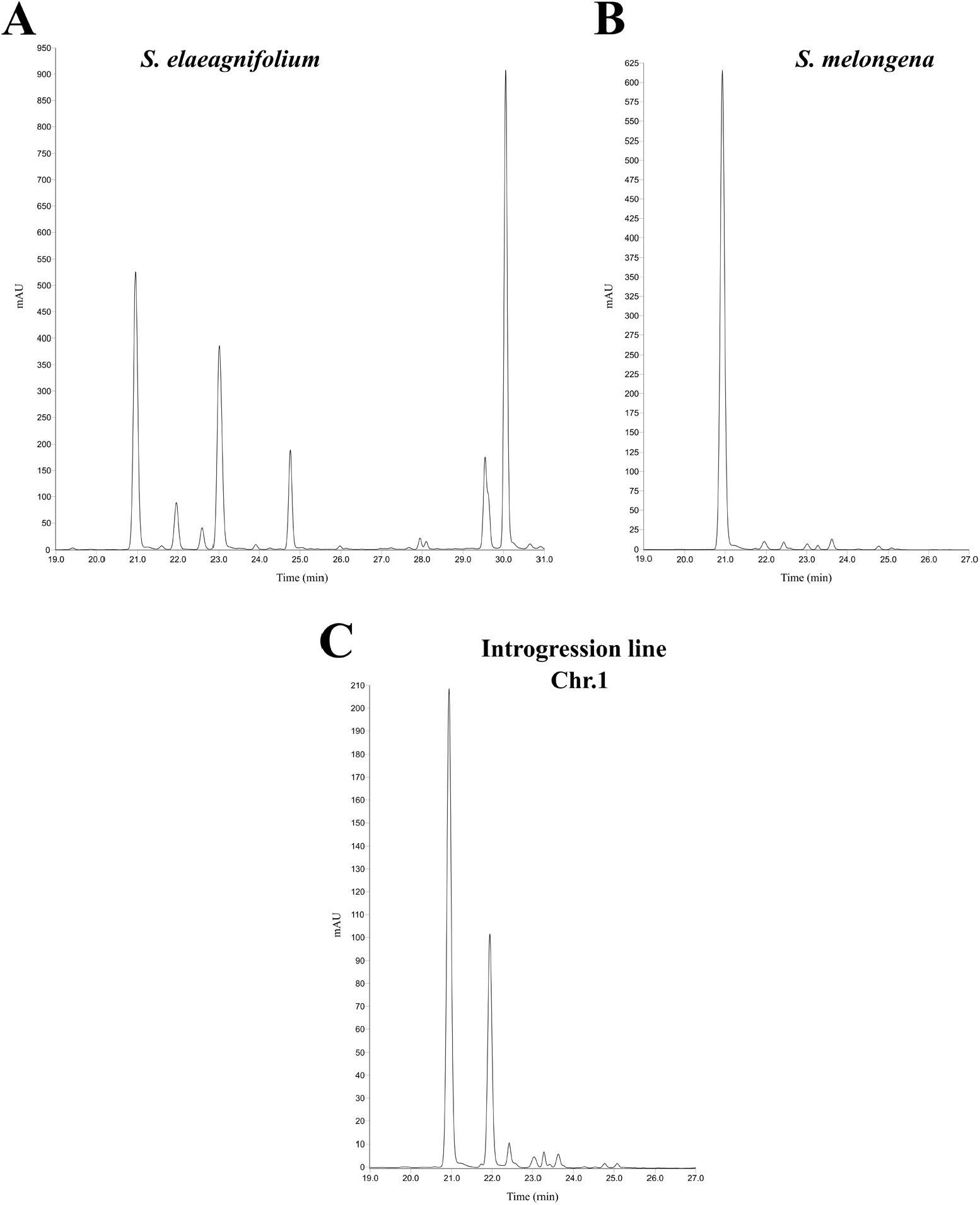
Representative HPLC chromatograms showing the caffeoylquinic acid (CQA) profiles of *S. melongena, S. elaeagnifolium*, and a representative backcross-derived recombinant line carrying an introgression from *S. elaeagnifolium* on chromosome 1. **(A)** *S. elaeagnifolium* (accession ELE2), **(B)** *S. melongena* (accession MEL3), and **(C)** a recombinant line of *S. melongena* carrying the *S. elaeagnifolium* introgression. Different y-axis scales are used for each panel.

### 3.2 Identification of the secondary peak as 4-caffeoylquinic acid

HPLC-MS/MS analysis of a representative eggplant extract from a plant carrying the chromosome 1 introgressed region and showing the differential profile revealed two distinct chromatographic peaks corresponding to caffeoylquinic acid (CQA) isomers (Figure 2A). First-dimension mass spectrometry (MS) showed that both compounds shared a parent ion at *m/z* 353.21 (Figure 2B), consistent with their identity as CQA isomers. The MS^2^ fragmentation pattern of the early-eluting major peak yielded a base peak at *m/z* 190.80, characteristic of the 5-CQA isomer (Figure 2C). In contrast, the MS^2^ spectrum of the late-eluting secondary peak showed a base peak at m/z 172.83, together with minor fragment ions at *m/z* 190.82 and *m/z* 134.86 (Figure 2D). According to the elution order and MS^2^ fragmentation criteria established by Clifford et al. (2003), these results support the identification of the late-eluting secondary peak as 4-caffeoylquinic acid (4-CQA).

**Figure 2.**
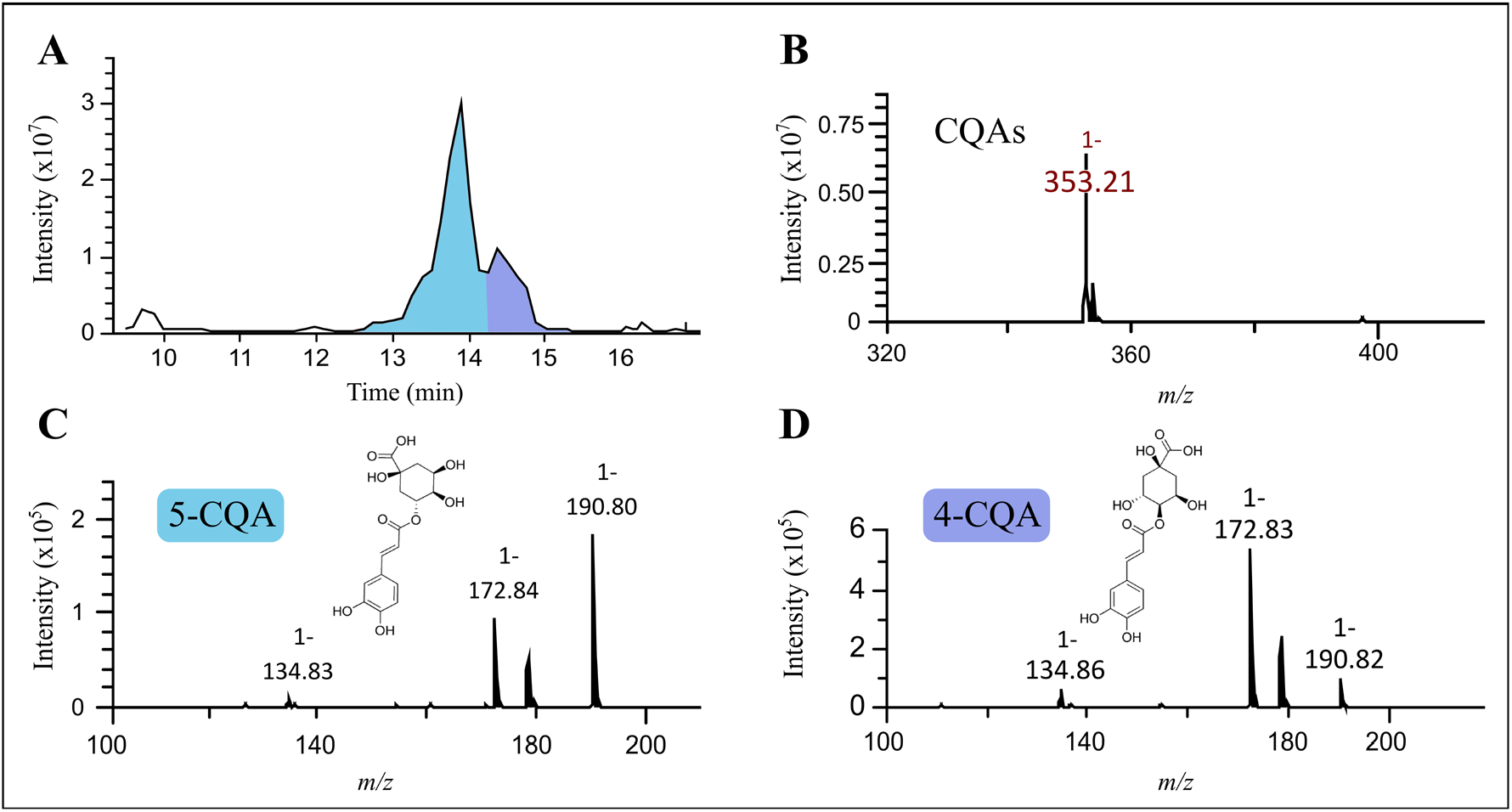
HPLC-MS/MS separation and identification of caffeoylquinic acid (CQA) isomers according to Clifford et al. (2003). **(A)** Chromatogram showing the elution profile of CQA isomers, with the major peak highlighted in light blue and the secondary peak in purple. **(B)** MS spectrum displaying the parent ion at *m/z* 353.21 for CQAs. **(C)** MS^2^ spectrum of the early-eluting major peak (light blue), identified as 5-CQA based on its base peak at *m/z* 190.80. **(D)** MS^2^ spectrum of the late-eluting secondary peak (purple), identified as 4-CQA, based on its base peak at *m/z* 172.83. Chemical structures for both isomers are provided in the insets.

### 3.3 Fine mapping of the chromosome 1 candidate region

Fine mapping of the chromosome 1 region associated with the secondary CQA peak was performed across three consecutive selfing generations. A total of 612 individuals were screened, yielding 96 recombinants (15.7%). SPET analysis confirmed 21 informative recombinants, corresponding to approximately 22% of the total recombinants across all generations, with 8 confirmed in the first generation (18.2%), 4 in the second (14.3%), and 9 in the third (37.5%). Combining these recombination breakpoints with the metabolic phenotype progressively reduced the candidate interval from 2.753 Mb (0.983-3.736 Mb) to 0.211 Mb (0.983-1.194 Mb), representing a 92.3% reduction of the initial region (Figure 3). Notably, the phenotypic evaluation of the segregating populations revealed that the secondary peak was present in both homozygous and heterozygous plants for the *S. elaeagnifolium* introgression, indicating a dominant genetic control for this trait. According to the annotation of the eggplant reference genome v5 (Gaccione et al. 2025), the final interval contains 30 annotated genes.

**Figure 3.**
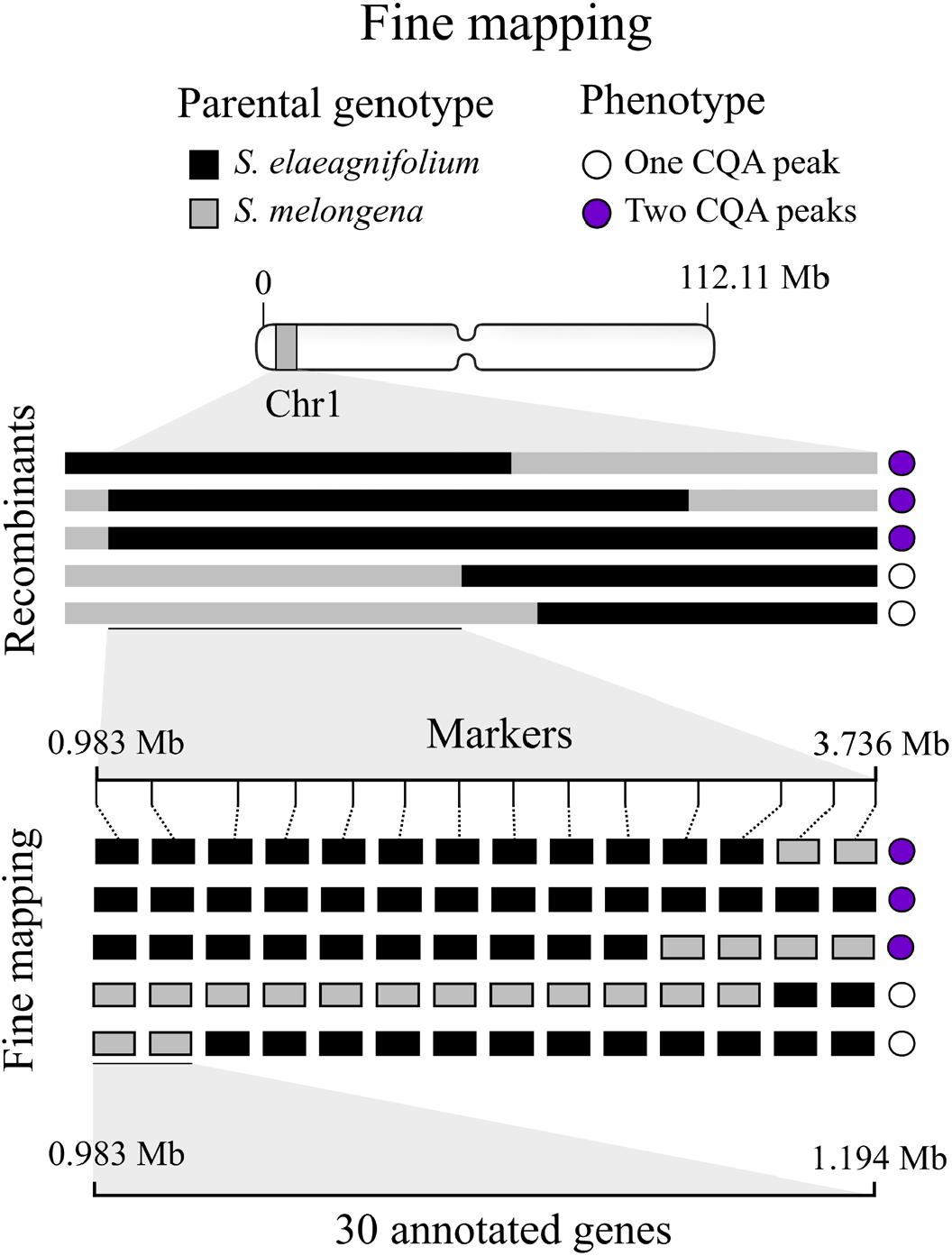
Fine mapping of the candidate region for the one vs. two CQA peak profile on chromosome 1 of eggplant. Recombinant lines derived from *S. melongena* (grey) and *S. elaeagnifolium* (black) were used to narrow the candidate region on chromosome 1. Phenotypes (purple: presence of two CQA peaks; white: presence of a single CQA peak) were associated with marker genotypes to delimit the critical region to 0.983-1.194 Mb, which contains 30 annotated genes.

### 3.4 Candidate gene analysis

Annotation of the 211 kb fine-mapped interval in the *S. melongena* reference genome identified 30 predicted genes (Table S2). The corresponding region in *S. elaeagnifolium* showed a comparable gene content (Table S3). Functional annotation of the predicted genes in both species revealed several shared gene classes across the interval, including endo-1,4-beta-xylanases, thaumatin-like proteins, an auxin transporter-like gene, a glycine-rich protein, translocase of chloroplast 120, dirigent proteins, amidases, a bHLH transcription factor, triosephosphate isomerase, acetylajmalan esterase-like genes, and an alkaline/neutral invertase (Tables S2 and S3).

At the structural level, microsynteny analysis showed that most of the candidate region was conserved between *S. melongena* and *S. elaeagnifolium*, with gene order and orientation broadly preserved across much of the interval (Figure 4). However, local structural differences were also detected, including genes for which no clear orthologous relationship could be established in the corresponding interval of the other species. The clearest divergence was observed in the region enclosed by the dashed box in Figure 4, corresponding to a structurally differentiated block between both genomes, including differences in gene number, gene length, intergenic distances, and orthology relationships.

**Figure 4.**
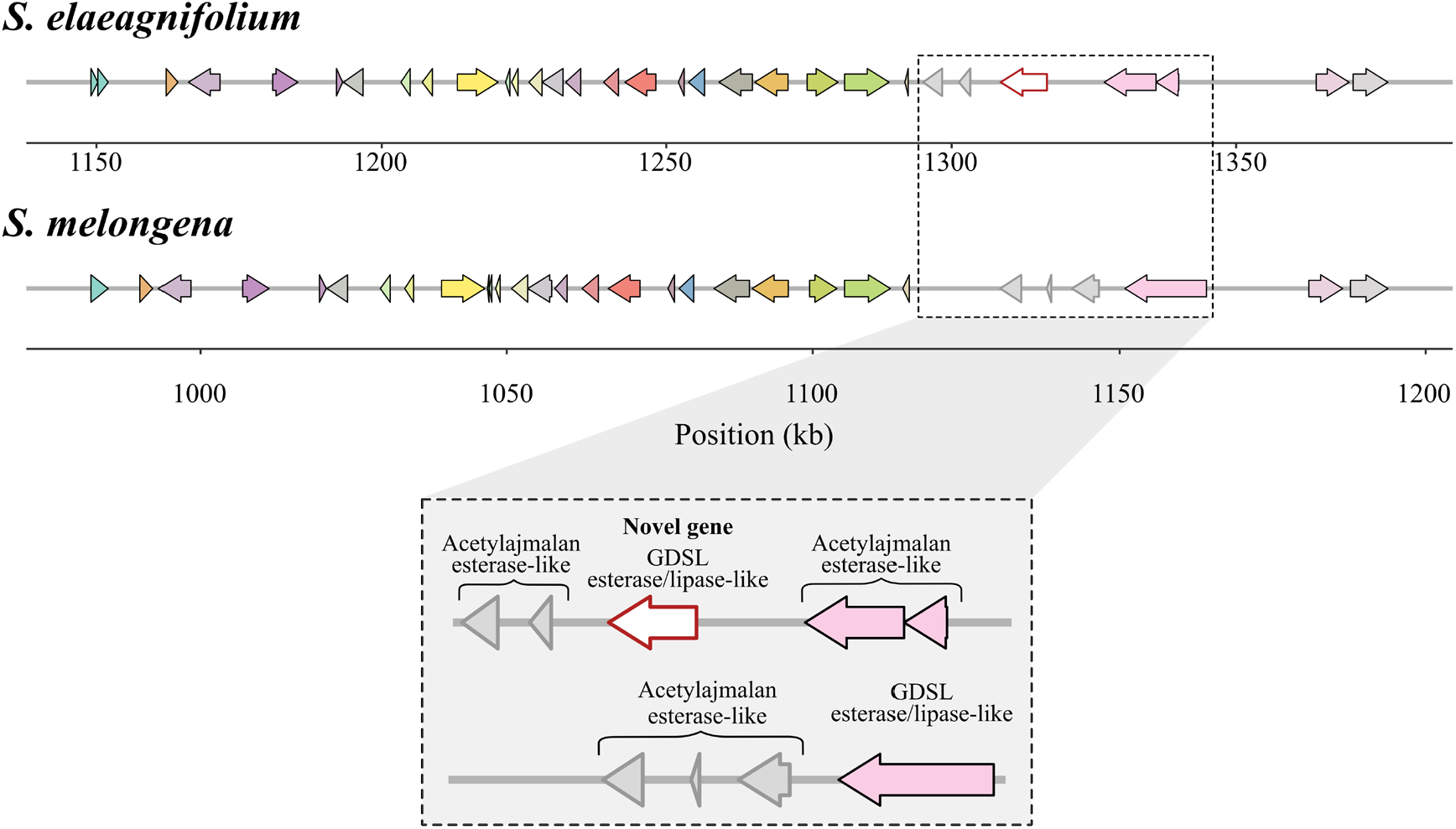
Structural microsynteny between *S. elaeagnifolium* (top) and *S. melongena* (bottom) for the fine-mapped candidate region associated with the differential CQA profile in chromosome 1. Arrows represent annotated genes, with arrow direction indicating gene orientation. Genes sharing the same color correspond to putative orthologs between the two species, whereas grey arrows indicate genes without an identified ortholog in the corresponding region. The enlarged box shows the organization of this differentiated block in both species and the predicted functional annotation of the genes located in this region, including acetylajmalan esterase-like/GDSL-type genes. The red-outlined arrow indicates a donor-specific novel gene identified in the *S. elaeagnifolium* interval through manual curation of public transcriptomic data. Genomic positions are shown in kb.

Within this structurally differentiated block, the *S. elaeagnifolium* interval contains a cluster of genes annotated as acetylajmalan esterase-like proteins, together with a transcript-supported novel gene identified by manual curation of public transcriptomic data (Figure 4). BLAST analysis of this novel gene revealed similarity to a GDSL esterase/lipase-like protein, providing a tentative functional annotation for this donor-specific gene model (Table S3). Taken together, the progressive reduction of the candidate interval through recombination and the identification of a localized structurally divergent block within this region point to this chromosome 1 segment as the main candidate region underlying the presence of the secondary 4-CQA peak. Although these results do not exclude the contribution of additional variants within this interval, they indicate that this region is a strong candidate to be the major determinant of the phenotype.

## 4. Discussion

By combining metabolite identification, fine mapping, and comparative microsynteny analysis, we delimited a candidate genomic interval associated with a secondary HPLC CQA-related peak detected in fruits of eggplant materials with a *S. elaeagnifolium* introgression in chromosome 1 (Villanueva et al. 2021). Fine mapping reduced the candidate region to 211 kb, and structural microsynteny analysis revealed a localized divergent block within this region despite overall conservation between both species. Together, these results associate the secondary 4-CQA peak with a small candidate interval showing local structural differences between *S. melongena* and *S. elaeagnifolium*.

The chlorogenic acid (CGA) pathway plays a central role in the phenolic profile of eggplant, with 5-caffeoylquinic acid (5-CQA) as the predominant compound in the fruit of both eggplant wild relatives and cultivated varieties (Plazas et al. 2013a; Rosa-Martínez et al. 2023a; Zhou et al. 2023). In contrast, 4-caffeoylquinic acid (4-CQA) is a less common isomer in solanaceous plants, typically found in minor quantities (Stommel and Whitaker 2003; Niño-Medina et al. 2017). However, its presence in significant amounts in *S. elaeagnifolium* derived eggplant materials represents a potentially novel source of metabolic variation in eggplant. The identification of 4-CQA as a secondary peak in these materials provides insight into the potential for modifying the metabolic profile of the eggplant fruit through introgressions from wild relatives. Previous studies have shown that caffeoylquinic acids (CQAs), including 4-CQA, possess antioxidant, anti-inflammatory and anticancer effects, highlighting the potential importance of this compound in enhancing the bioactive properties of eggplant (Chen et al. 2025). Furthermore, understanding the structural basis of CGA biosynthesis and the ability to modulate CQA levels in crop breeding could offer new strategies for improving functional food properties or for developing plants with specific metabolic profiles (Mennella et al. 2010; Lallemand et al. 2012; Docimo et al. 2016).

Wild relatives, such as *S. elaeagnifolium*, are recognized as valuable sources of metabolic diversity, which can be introgressed into cultivated species to broaden their phenolic profiles and enhance their functional properties, offering new opportunities to improve both the nutritional and bioactive value of eggplant (Kaushik et al. 2015; Rosa-Martínez et al. 2022; Kozuharova et al. 2024). In this regards, *S. elaeagnifolium* presents a much more complex phenolic profile than *S. melongena*, characterized by a broader diversity of phenolic compounds in the fruit, which is reflected in its chromatographic profile. The presence of multiple peaks, including the secondary peak corresponding to 4-CQA, aligns with the complex metabolic profile of *S. elaeagnifolium* observed in previous studies (Al-hamaideh et al. 2020; Kobisi et al. 2024). These studies have demonstrated the presence of key phenolic compounds s, such as quercetin, gallic acid, kaempferol, and naringenin, all of which contribute to the plant’s antioxidant and antiinflammatory activities, while also highlighting their potential as source of antidiabetic and antimicrobial compounds (Bouslamti et al. 2022, 2023; Francis Xavier et al. 2025).

The introgressions of *S. elaeagnifolium* into cultivated eggplant (*S. melongena*) are of particular interest because they involve the transfer of genomic segments from a highly divergent tertiary genepool relative (Syfert et al. 2016). This evolutionary and structural divergence between the two species (Särkinen et al. 2013; Knapp et al. 2017) may underlie the metabolic differences observed here, supporting that *S. elaeagnifolium* can contribute novel metabolic traits to the eggplant genome through introgression.

The characterization of the genetic basis underlying fruit quality has been a major goal in eggplant breeding (Taher et al. 2017). Some efforts have successfully developed linkage maps to identify QTLs associated with key biochemical traits, including chlorogenic acid and other metabolic compounds (Toppino et al. 2016; Rosa-Martínez et al. 2023a; Gaccione et al. 2025). In this context, fine mapping of the candidate region on chromosome 1, based on the *S. melongena* reference genome, reduced the interval to a 211 kb segment containing 30 annotated genes. This region was precisely defined by combining phenotypic data with high-throughput genotyping of recombinant lines, enabling the identification of key recombination events. Our fine mapping builds upon previous genome-wide association efforts by Villanueva et al. (2021), who first identified a highly significant, previously unreported QTL in a broader region of chromosome 1 in their study of advanced backcrosses of *S. elaeagnifolium* into *S. melongena*. The same differential pattern was detected by Villanueva et al. (2021) in plants grown in soil under open-field conditions, whereas in our study it was consistently observed across consecutive years in introgression materials evaluated under greenhouse conditions. Given that CQA levels can be modulated by environmental factors such as location, cultivation system, light, temperature, and drought (Stommel et al. 2015; Chen et al. 2025), the recurrence of this secondary peak across different years supports the stability of the trait across the tested conditions. Furthermore, its expression in both homozygous and heterozygous introgression-derived materials indicates a dominant control of the trait, showing that one copy of the introgressed *S. elaeagnifolium* segment is sufficient to generate the secondary 4-CQA peak. Together with the localized structural divergence identified in this interval, this observation supports the idea that the wild haplotype carries a functional determinant that is missing or functionally reduced in the corresponding *S. melongena* region.

The reduced candidate region identified on chromosome 1 encompasses several genes that may play crucial roles in the biosynthesis or regulation of CQA-related compounds. Structural microsynteny analysis of this interval revealed a localized divergent block, indicating that despite the overall genomic conservation between *S. melongena* and *S. elaeagnifolium*, significant local structural differences exist (Benoit et al. 2025; Gaccione et al. 2025). Such localized structural variations (SVs) are increasingly recognized as primary drivers of phenotypic and metabolic diversity in crops, including eggplant (Gao et al. 2019; Gaccione et al. 2025). This specific divergence strongly supports the idea that the wild introgression introduces novel genetic elements directly responsible for the altered metabolite profiles, particularly the presence of the secondary 4-CQA peak (Sulli et al. 2021; Villanueva et al. 2021; Rosa-Martínez et al. 2023b).

Focusing on the genetic composition of this structurally divergent block, we found that the localized genes are notably annotated as members of the GDSL lipase and acetylajmalan esterase families, including a transcript-supported putative novel gene identified from transcriptomic data. Given that acetylajmalan esterases are formally classified as a subclass within the broader GDSL lipase/esterase superfamily, this locus essentially comprises a cluster of GDSL-type enzymes (Ruppert et al. 2005). These enzymes are known for their vast functional diversity, with proven roles in modifying phenolic compounds, lipid metabolism, plant defence, and tolerance to stress (Tan et al. 2014; Ding et al. 2019; Tang et al. 2025). More specifically, previous studies in other Solanaceae species have demonstrated that GDSL enzymes are actively involved in phenylpropanoid and chlorogenic acid metabolism, including the hydrolysis and synthesis of di-CQA (Teutschbein et al. 2010; Miguel et al. 2020; Guo et al. 2025). Given this functional precedent, it is biochemically plausible that the enzymes encoded by this introgressed GDSL-like gene cluster facilitate a specific molecular rearrangement, such as driving the isomerization of quinic acid derivatives or preferentially releasing 4-CQA from more complex precursors. Consequently, this introgressed cluster of genes emerges as the primary candidate locus underlying the novel 4-CQA phenotype. From a breeding perspective (Prohens et al. 2017), delimiting this region and its associated markers provides a practical tool for tracking the introgression and selecting eggplant lines with a more diverse CQA profile, thereby facilitating the development of materials with more diverse and potentially enhanced functional quality.

## Conclusions

The present work provides a more precise characterization of the distinct caffeoylquinic acid profile associated with a *Solanum elaeagnifolium* introgression on chromosome 1 in eggplant. The secondary CQA-related peak was identified as 4-*O*-caffeoylquinic acid (4-CQA), resolving the identity of a previously uncharacterized compound associated with these introgression materials. Fine mapping reduced the associated interval from 2.753 Mb to 0.211 Mb, delimiting a small genomic region linked to this phenotype. Comparative microsynteny analysis revealed a localized structurally divergent block between *S. melongena* and *S. elaeagnifolium*. Within this block, a cluster of acetylajmalan esterase-like/GDSL-type genes was identified, including a transcript-supported putative novel donor-specific gene, and these emerged as particularly relevant candidates associated with the distinct CQA profile. Together, these results support the value of *S. elaeagnifolium* introgressions as a source of phenolic diversification in eggplant and provide useful markers for tracking this region in future breeding programmes aimed at developing lines with a more diverse CQA profile and potentially improved functional properties.

## Acknowledgements

This work was funded by the Universitat Politècnica de València through project PAID-06-25, by the Spanish Ministerio de Ciencia, Innovación y Universidades through MICIU/AEI/10.13039/501100011033 and ERDF/EU (grant PID2024-160953OB-I00), by MICIU/AEI/10.13039/501100011033 and co-funded by the European Union (grant PDC2025-165434-I00); and by the Conselleria d’Educació, Cultura, Universitats i Ocupació of the Generalitat Valenciana (grant CIPROM/2021/020). GV also acknowledges the Universitat Politècnica de València (UPV) for a postdoctoral grant (PAID-10-24). PG has received a postdoctoral grant (RYC2021-031999-I) funded by MICIU/AEI/10.13039/501100011033 and by the European Union NextGeneration EU/PRTR. VB-F has received a predoctoral grant (CIACIF/2023/238) funded by Conselleria d’Educació, Cultura, Universitats i Ocupació (Generalitat Valenciana).

