## Supplemental Tables for "Genetic dissection of a *Solanum elaeagnifolium* tertiary genepool introgression associated with 4-*O*-caffeoylquinic acid accumulation in eggplant"

Table S1. Primers used to target candidate genomic regions based on the eggplant genome v5, including their sequences, amplicon sizes, and melting temperatures.

| Target region | Primer Name | Sequence (5' → 3') | Amplicon size (bp) | Tm (ºC) |
|---|---|---|---|---|
| Chr1:2,393,500 (V5) | SET1_5_14_F | CCCATCCAAACACAGGCAAT | 155 | 58 |
| | SET1_5_14_R | TGTTCCACCTGCAAAACGAG | | 58 |
| Chr1:2,451,000 (V5) | SET1_5_21_F | TGGTGGCTTGTCGTTTTAGC | 102 | 58 |
| | SET1_5_21_R | ACCCCAGAGAGATTTGAAAGC | | 60 |
| Chr1:3,141,300 (V5) | SET1_5_94_F | CAGAACGGAAAGTGAAGAACAGT | 100 | 60 |
| | SET1_5_94_R | GGAATCCAAGTCGTCAAGGG | | 62 |
| Chr1:3,657,600 (V5) | SET1_6_5_F | ACTCATCCCAACCACTACCTG | 118 | 60 |
| | SET1_6_5_R | GAGGGAGACTGCTTGGAGTT | | 60 |
| Chr1:1,880,900 (V5) | SET2_4.58_F | CGGAACCTAACCTAAATTCTGGG | 100 | 63 |
| | SET2_4.58_R | ACTGATCAGCGCTCTCACC | | 59 |
| Chr1:2,393,500 (V5) | SET2_5.15_F | CGCAAAGCAAGATCCCACAA | 155 | 58 |
| | SET2_5.15_R | TGTTGGTTAGCTTTGTGGCC | | 58 |
| Chr1:2,814,800 (V5) | SET2_5.59_F | TCCCTCATTCTGGTTGGAGC | 126 | 60 |
| | SET2_5.59_R | GCACTCGGATCAAAACCTGT | | 58 |
| Chr1:1,262,000 (V5) | SET3_1.35_F | CGTCTCTCCTACCCTTTGCA | 130 | 60 |
| | SET3_1.35_R | AGGGATAGAGATTATAGAAGAG | | 57 |
| Chr1:1,513,000 (V5) | SET3_1.62_F | TCAAGAGGCCAATCTGAAGGA | 100 | 59 |
| | SET3_1.62_R | TACGTCAGGATCAGGAGGGA | | 60 |
| Chr1:1,740,600 (V5) | SET3_1.87_F | CTTCACTGTAGTCACCCAAAAC | 101 | 60 |
| | SET3_1.87_R | GATGAGATAAGGTAGCTTATGC | | 58 |
| Chr1:1,785,000 (V5) | SET3_1.92_F | TCATGGCTTCATTCTTCGACA | 95 | 57 |
| | SET3_1.92_R | CCGAAACTAGATCATCACACTTT | | 59 |
| Chr1:2,476,800 (V5) | SET3_2.68_F | AGTCTTGAATCTCTCTGCGGT | 197 | 59 |
| | SET3_2.68_R | TGCCCTTTGACTGGTATGGA | | 58 |
| Chr1:2,696,300 (V5) | SET3_2.91_F | AGAACTACAAGTTAATTATCTGTGG | 64 | 59 |
| | SET3_2.91_R | TAGGTAATTAGTACCTTTCACTGG | | 60 |

Table S2. Annotated candidate genes identified within the 211 kb fine-mapped candidate interval on chromosome 1 of *Solanum melongena* ('67/3', v5).

| Gene ID | Chromosome | Position (Start-End, bp) | Length (kb) | Annotated Function |
|---|---|---|---|---|
| SMEL5_01g001230 | 1 | 982238-985044 | 2,806 | similar to XP_049388910.1: endo-1,4-beta-xylanase 5 isoform X1 (*Solanum stenotomum* ) |
| SMEL5_01g001240 | 1 | 990215-992304 | 2,089 | similar to XP_055815554.1: thaumating-like protein 1 isoform X1 (*Solanum dulcamara* ) |
| SMEL5_01g001250 | 1 | 993198-998427 | 5,229 | similar to XP_055815557.1: thaumating-like protein 1b (*Solanum dulcamara* ) |
| SMEL5_01g001260 | 1 | 1010562-1011188 | 0,626 | similar to XP_009621128.1: auxin transporter-like protein 2 (*Nicotiana tomentosiformis* ) |
| SMEL5_01g001270 | 1 | 1019416-1020520 | 1,104 | similar to XP_016539926.1: glycine-rich protein 2-like (*Capsicum annuum* ) |
| SMEL5_01g001280 | 1 | 1020655-1024011 | 3,356 | similar to PSS21742.1: Vacuolar protein (*Actinidia chinensis* var.*chinensis* ) |
| SMEL5_01g001290 | 1 | 1029359-1030971 | 1,612 | similar to XP_015170007.1: PREDICTED: PI-PLC X-box domain-containing protein DDB_GO293739-like isoform X2 (*Solanum tuberosum* ) |
| SMEL5_01g001300 | 1 | 1033353-1034803 | 1,45 | similar to XP_006359666.1: PREDICTED: PI-PLC X-box domain-containing protein DDB_GO293730-like (*Solanum tuberosum* ) |
| SMEL5_01g001310 | 1 | 1039356-1046436 | 7,08 | similar to XP_049350615.1: translocase of chloroplast 120, chloroplastic-like (*Solanum verrucosum* ) |
| SMEL5_01g001320 | 1 | 1046867-1047117 | 0,25 | similar to XP_055814722.1: dirigent protein 1-like (*Solanum dulcamara* ) |
| SMEL5_01g001330 | 1 | 1047188-1047575 | 0,387 | similar to XP_060169332.1: dirigent protein 1-like (*Lycium barbarum* ) |
| SMEL5_01g001340 | 1 | 1048206-1048936 | 0,73 | similar to XP_055814723.1: dirigent protein 23-like (*Solanum dulcamara* ) |
| SMEL5_01g001350 | 1 | 1050788-1053447 | 2,659 | similar to XP_049350624.1: probable amidase at4g34880 (*Solanum verrucosum* ) |
| SMEL5_01g001360 | 1 | 1053448-1057353 | 3,905 | similar to XP_027767518.1: probable amidase at4g34880 (*Solanum pennellii* ) |
| SMEL5_01g001370 | 1 | 1057880-1059861 | 1,981 | similar to XP_060168583.1: protodermal factor 1 (*Lycium barbarum* ) |
| SMEL5_01g001380 | 1 | 1063596-1064989 | 1,393 | similar to XP_055815563.1: eukaryotic translation initiation factor 4B3 (*Solanum dulcamara* ) |
| SMEL5_01g001390 | 1 | 1066581-1071785 | 5,204 | similar to XP_006359660.1: PREDICTED: probable magnesium transporter NIPA6 isoform X1(*Solanum tuberosum* ) |
| SMEL5_01g001400 | 1 | 1076355-1077419 | 1,064 | similar to XP_055815566.1: peptidyl-prolyl cis-trans isomerase (*Solanum dulcamara* ) |
| SMEL5_01g001410 | 1 | 1078173-1079912 | 1,739 | similar to XP_047251154.1: proline-rich protein 4 isoform X2 (*Capsicum annuum* ) |
| SMEL5_01g001420 | 1 | 1083867-1089661 | 5,794 | similar to XP_049378090.1: translocon-associated protein subunit alpha-like (*Solanum stenotomum* ) |
| SMEL5_01g001430 | 1 | 1090098-1096004 | 5,906 | similar to XP_006359652.1: PREDICTED: protein XAP5 CIRCADIAN TIMEKEEPER (*Solanum tuberosum* ) |
| SMEL5_01g001440 | 1 | 1099470-1103961 | 4,491 | similar to XP_055815571.1: transcription factor bHLH121 isoform X1 (*Solanum dulcamara* ) |
| SMEL5_01g001450 | 1 | 1109856-1109941 | 0,085 | similar to XP_049387904.1: triosephosphate isomerase, chloroplastic-like (*Solanum stenotomum* ) |
| SMEL5_01g001460 | 1 | 1114572-1115636 | 1,064 | similar to XP_007131593.1: hypothetical protein PH AVU_011G026300g (*Phaseolus vulgaris* ) |
| SMEL5_01g001470 | 1 | 1130483-1133915 | 3,432 | similar to XP_055816753.1: acetylajmalan esterase-like isoform X1 (*Solanum dulcamara* ) |
| SMEL5_01g001480 | 1 | 1138473-1138680 | 0,207 | similar to XP_055816753.1: acetylajmalan esterase-like isoform X1 (*Solanum dulcamara* ) |
| SMEL5_01g001490 | 1 | 1142196-1146621 | 4,425 | similar to XP_055816753.1: acetylajmalan esterase-like isoform X1 (*Solanum dulcamara* ) |
| SMEL5_01g001500 | 1 | 1150845-1164186 | 13,341 | similar to XP_055816754.1: GDSL esterase/lipase At5g03980-like isoform X2 (*Solanum dulcamara* ) |
| SMEL5_01g001510 | 1 | 1184727-1185286 | 0,559 | similar to XP_059314339.1: probable alkaline/neutral invertase B (*Lycium ferocissimum* ) |
| SMEL5_01g001520 | 1 | 1187701-1193827 | 6,126 | similar to XP_015062990.1: putative ER lumen protein-retaining receptor C28H8.4 (*Solanum pennellii* ) |

Table S3. Annotated candidate genes identified in the syntenic interval of *Solanum elaeagnifolium* corresponding to the fine-mapped candidate region on chromosome 1.

| Gene ID | Chromosome | Position (Start-End, bp) | Length (kb) | Annotated Function |
|---|---|---|---|---|
| Solanum_eleagnifolium_Selea_Chr01_000061.1 | 1 | 1149047-1150240 | 1,194 | similar to XP_006359674.2: endo-1,4-beta-xylanase UM03411 isoform X1 (*Solanum tuberosum* ) |
| Solanum_eleagnifolium_Selea_Chr01_000062.1 | 1 | 1150243-1152067 | 1,825 | similar to XP_055815552.1: endo-1,4-beta-xylanase 5-like isoform X1 (*Solanum dulcamara* ) |
| Solanum_eleagnifolium_Selea_Chr01_000063.1 | 1 | 1162097-1164182 | 2,086 | similar to XP_055815554.1: thaumatin-like protein 1 isoform X1 (*Solanum dulcamara* ) |
| Solanum_eleagnifolium_Selea_Chr01_003838.1 | 1 | 1166005-1171567 | 5,563 | similar to XP_055815556.1: thaumatin-like protein 1b (*Solanum dulcamara* ) |
| Solanum_eleagnifolium_Selea_Chr01_000064.1 | 1 | 1180814-1185264 | 4,451 | similar to XP_015069200.1: auxin transporter-like protein 5 (*Solanum pennellii* ) |
| Solanum_eleagnifolium_Selea_Chr01_000065.1 | 1 | 1192028-1193107 | 1,08 | similar to XP_055815561.1: glycine-rich protein 2-like (*Solanum dulcamara* ) |
| Solanum_eleagnifolium_Selea_Chr01_003837.1 | 1 | 1193266-1196716 | 3,451 | similar to XP_049377735.1: uncharacterized protein LOC125842474 (*Solanum stenotomum* ) |
| Solanum_eleagnifolium_Selea_Chr01_003836.1 | 1 | 1203360-1204980 | 1,621 | similar to XP_049350633.1: uncharacterized protein LOC125815242 (*Solanum verrucosum* ) |
| Solanum_eleagnifolium_Selea_Chr01_003835.1 | 1 | 1207116-1208906 | 1,791 | similar to XP_049350634.1: uncharacterized protein LOC125815243 (*Solanum verrucosum* ) |
| Solanum_eleagnifolium_Selea_Chr01_000066.1 | 1 | 1213278-1220468 | 7,191 | similar to XP_006359664.1: translocase of chloroplast 120, chloroplastic (*Solanum tuberosum* ) |
| Solanum_eleagnifolium_Selea_Chr01_003834.1 | 1 | 1221765-1222514 | 0,75 | similar to XP_006359663.1: dirigent protein 1-like (*Solanum tuberosum* ) |
| Solanum_eleagnifolium_Selea_Chr01_003833.1 | 1 | 1222840-1223934 | 1,095 | similar to XP_055814723.1: dirigent protein 23-like (*Solanum dulcamara* ) |
| Solanum_eleagnifolium_Selea_Chr01_003832.1 | 1 | 1225851-1228170 | 2,32 | similar to XP_049350624.1: probable amidase At4g34880 (*Solanum verrucosum* ) |
| Solanum_eleagnifolium_Selea_Chr01_003831.1 | 1 | 1228172-1231836 | 3,665 | similar to XP_027767518.1: probable amidase At4g34880 (*Solanum pennellii* ) |
| Solanum_eleagnifolium_Selea_Chr01_003830.1 | 1 | 1232317-1234959 | 2,643 | similar to XP_006359661.1: uncharacterized protein (*Solanum tuberosum* ) |
| Solanum_eleagnifolium_Selea_Chr01_003829.1 | 1 | 1238977-1241601 | 2,625 | similar to XP_055815563.1: eukaryotic translation initiation factor 4B3 (*Solanum dulcamara* ) |
| Solanum_eleagnifolium_Selea_Chr01_003828.1 | 1 | 1242699-1248169 | 5,471 | similar to XP_016539910.2: probable magnesium transporter NIPA6 (*Capsicum annuum* ) |
| Solanum_eleagnifolium_Selea_Chr01_003827.1 | 1 | 1252129-1253132 | 1,004 | similar to XP_055815566.1: peptidyl-prolyl cis-trans isomerase (*Solanum dulcamara* ) |
| Solanum_eleagnifolium_Selea_Chr01_003826.1 | 1 | 1253894-1256718 | 2,825 | similar to XP_047251154.1: proline-rich protein 4 isoform X2 (*Capsicum annuum* ) |
| Solanum_eleagnifolium_Selea_Chr01_003825.1 | 1 | 1259244-1265136 | 5,893 | similar to XP_049378090.1: translocon-associated protein subunit alpha-like (*Solanum stenotomum* ) |
| Solanum_eleagnifolium_Selea_Chr01_003824.1 | 1 | 1265531-1271390 | 5,86 | similar to XP_004230880.1: protein XAP5 CIRCADIAN TIMEKEEPER (*Solanum lycopersicum* ) |
| Solanum_eleagnifolium_Selea_Chr01_000067.1 | 1 | 1274746-1280193 | 5,448 | similar to XP_019067474.1: transcription factor bHLH121 isoform X1 (*Solanum lycopersicum* ) |
| Solanum_eleagnifolium_Selea_Chr01_000068.1 | 1 | 1281325-1289165 | 7,841 | similar to XP_006359648.1: triosephosphate isomerase, chloroplastic (*Solanum tuberosum* ) |
| Solanum_eleagnifolium_Selea_Chr01_003823.1 | 1 | 1291785-1292372 | 0,588 | similar to XP_004230988.1: uncharacterized protein (*Solanum lycopersicum* ) |
| Solanum_eleagnifolium_Selea_Chr01_003822.1 | 1 | 1295002-1298326 | 3,325 | similar to XP_055816753.1: acetylajmalan esterase-like isoform X1 (*Solanum dulcamara* ) |
| Solanum_eleagnifolium_Selea_Chr01_003821.1 | 1 | 1301290-1303276 | 1,987 | similar to XP_055816747.1: acetylajmalan esterase-like isoform X1 (*Solanum dulcamara* ) |
| Novel_gene_Chr01_candidate_region | 1 | 1308603-1316748 | 8,146 | similar to XP_049407169.1:GDSL esterase/lipase At5g03980-like (*Solanum stenotomum* ) |
| Solanum_eleagnifolium_Selea_Chr01_003820.1 | 1 | 1326814-1335923 | 9,11 | similar to XP_055816747.1: acetylajmalan esterase-like isoform X1 (*Solanum dulcamara* ) |
| Solanum_eleagnifolium_Selea_Chr01_003819.1 | 1 | 1335925-1339900 | 3,976 | similar to XP_055816753.1: acetylajmalan esterase-like isoform X1 (*Solanum dulcamara* ) |
| Solanum_eleagnifolium_Selea_Chr01_000069.1 | 1 | 1364078-1369860 | 5,783 | similar to XP_059314337.1: probable alkaline/neutral invertase B (*Lycium ferocissimum* ) |
| Solanum_eleagnifolium_Selea_Chr01_000070.1 | 1 | 1370543-1376676 | 6,134 | similar to XP_004230929.1: uncharacterized protein (*Solanum lycopersicum* ) |
